# The power of language: functional brain network topology of deaf and hearing in relation to sign language experience

**DOI:** 10.1101/335414

**Authors:** Michel R.T. Sinke, Jan W. Buitenhuis, Frank van der Maas, Job Nwiboko, Rick M. Dijkhuizen, Eric van Diessen, Willem M. Otte

**Author notes:** Shared last author (Eric van Diessen, Willem M. Otte). Corresponding author: Michel R.T. Sinke, MSc Biomedical MR Imaging and Spectroscopy group, Center for Image Sciences, University Medical Center Utrecht, Heidelberglaan 100, 3584 CX Utrecht, The Netherlands.; ORCID ID: https://orcid.org/0000-0002-8185-9209.

## Abstract

Prolonged auditory sensory deprivation leads to brain reorganization, indicated by functional enhancement in remaining sensory systems, a phenomenon known as cross-modal plasticity. In this study we investigated differences in functional brain network shifts from eyes-closed to eyes-open conditions between deaf and hearing people. Electroencephalography activity was recorded in deaf (N = 71) and hearing people (N = 122) living in rural Africa, which yielded a unique data-set of congenital, pre-lingual and post-lingual deaf people, with a divergent experience in American Sign Language. Functional networks were determined from the synchronization of electroencephalography signals between fourteen electrodes distributed over the scalp. We studied the synchronization between the auditory and visual cortex and performed whole-brain minimum spanning tree analysis based on the phase lag index of functional connectivity. This tree analysis accounts for variations in global network density and allows unbiased characterization of functional network backbones. We found increased functional connectivity between the auditory and visual cortex in deaf people during the eyes-closed condition in both the alpha and beta bands. Furthermore, we found functional network backbone shifts both in deaf and healthy people as they went from eyes-closed to eyes-open conditions. In both the alpha and beta band the deafs’ brain showed larger functional backbone-shifts in node strength compared to controls. In the alpha band this shift in network strength differed among deaf participants and depended on type of deafness: congenital, pre-lingual or post-lingual deafness. In addition, a correlation was found between functional backbone characteristics and experience of sign language. Our study revealed more insights in functional network reorganization specifically due to prolonged lack of auditory input, but might also be helpful for sensory deprivation and cross-modal plasticity in general. Global cortical network reorganization in deaf people supports the plastic capacities of the young brain. The differences between type of deafness stresses that etiology affects functional reorganization, whereas the association between network organization and acquired sign language experience reflects ongoing brain adaptation in people with hearing disabilities.

## Introduction

Audition is an important sensory modality, which is essential for daily life and communication. Impairment or loss of hearing interferes with many activities, specifically limiting communication with others, which could easily lead to social isolation. The prevalence of this serious disability is greatest in middle- and low-income countries (Durkin, 2002; Stevens et al., 2013; WHO, 2014). While in the United States about two out of every 1,000 children is born with disabling hearing loss (Vohr, 2003), this number is considerably higher in Sub-Saharan Africa where about two percent of the children is born with disabling hearing loss (WHO, 2012). A subset of these children has profound hearing loss, resulting in absolute deafness. Infectious diseases are a major cause of deafness in these regions (Mulwafu et al., 2016).

Prolonged periods of deafness are believed to cause profound neural reorganization related to functional enhancement in remaining sensory systems, which is referred to as cross-modal plasticity (Bavelier and Neville, 2002; Merabet and Pascual-Leone, 2010; Ptito et al., 2001). Deaf people have to rely more on visual input (and input from other modalities), which consequently leads to increased involvement of neuronal networks processing visual information (Bavelier et al., 2006; Bosworth and Dobkins, 2002; Brozinsky and Bavelier, 2004; Dye et al., 2007; Finney and Dobkins, 2001; Hauser et al., 2007). While this reorganization occurs inevitably as a result of profound deafness, cross-modal plasticity is also strongly related to the acquisition and use of sign language and lip reading (Meyer et al., 2007; Pénicaud et al., 2013). Furthermore, the extent of cross-modal plasticity is also dependent on the age of onset and the duration of deafness (Brotherton et al., 2016; Li et al., 2013), as well as on the use of hearing aid and degree of auditory deprivation (Doucet et al., 2006; Shiell et al., 2014). Cross-modal plasticity has been reported in different studies on deaf as well as blind people (Collignon et al., 2011; Klinge et al., 2010; Lewis et al., 2010; Liu et al., 2007; Ptito and Kupers, 2005; Renier et al., 2014; Théoret et al., 2004; Yu et al., 2008).

Brain mapping techniques such as functional magnetic resonance imaging (MRI) and electroencephalography (EEG) enable detection of functional neural reorganization in deaf (as compared to hearing people), as well as assessment of differences between early-deafness and late-deafness, e.g. by using task-based or resting-state paradigms. For example it has been shown that cortical auditory or auditory association areas in deaf people are responsive to visual motion stimuli. These auditory regions include the planum temporale (Petitto, 2000; Sadato et al., 2005; Shiell et al., 2016) and primary auditory cortices, like posterior superior temporal gyrus (Almeida et al., 2015; Ding et al., 2015; Karns et al., 2012; Li et al., 2015) and Hechl’s gyrus (Karns et al., 2012; Meyer et al., 2007; Scott et al., 2014; Smith et al., 2011). Furthermore, functional MRI studies have shown that the middle superior temporal sulcus is more prominently activated by visual stimuli in early-deaf subjects than in late-deaf subjects (Li et al., 2013; Neville et al., 1998; Sadato et al., 2004). Although these task-based approaches have provided valuable insights in cross-modal plasticity, they do not take into account the mutual dependency of different functional regions and the integrative nature of the human brain to process auditory information (Hackett, 2012).

The human brain forms a complex integrative network, which consists of spatially distributed, but functionally connected (i.e. synchronized activated) regions that continuously interact with each other (Bullmore and Sporns, 2009; van den Heuvel and Hulshoff Pol, 2010). Brain function and reorganization can only be properly understood when studied in their context, i.e. within a functional network, which can be mapped by electroencephalography or functional MRI (Stam and van Straaten, 2012). A powerful approach to assess organizational aspects of functional brain networks is provided by graph analysis (Bullmore and Sporns, 2012). Graph analysis describes a complex system like the human brain, as a set of nodes (i.e. functional brain regions such as the auditory or visual cortex) and edges or ties (i.e. the functional connections between regions), and provides quantitative information on the topological properties of networks (Bullmore and Sporns, 2009; Heuvel et al., 2012; Rubinov and Sporns, 2010). It has been shown that the healthy human brain can be characterized as a complex network that effectively combines integration (i.e. global efficiency, indicated by a short average path length) and segregation (i.e. functionally specialized brain regions (Bullmore and Sporns, 2012), indicated by a high clustering coefficient), which together is defined as a small-world organization (Bullmore and Sporns, 2009; Watts and Strogatz, 1998). Other important aspects of brain organization, typified by graph analyses, are modularity (i.e. different functional modules) and hubness (i.e. relatively small number of highly connected nodes) (Barabasi and Albert, 1999).

So far only a few studies have used graph analysis to examine structural or functional brain networks in deaf subjects. Kim et al. showed that pre-lingual deaf adults have altered morphological networks compared to normal controls, whereas post-lingual deaf adults did not show differences with normal controls, indicating that auditory experience might affect the morphology of brain networks in deaf adults (Kim et al., 2014). Li et al. (2016) showed increased connectivity between the limbic system and regions involved in visual and language processing, as well as decreased connectivity between the visual and language processing regions. However small-worldness was not changed in pre-lingual deaf adults as compared to hearing controls (Li et al., 2016).

Although classical graph analysis has revealed significant aspects of brain reorganization in deaf people, and despite its potential and popularity (Giusti et al., 2016), it has some intrinsic limitations, particularly for inter-subject or between-group comparisons where network sizes and densities are different (van Wijk et al., 2010). Commonly used network metrics, such as the clustering coefficient (i.e. segregation) and average path length (i.e. integration or global efficiency) are highly affected by the number of connections (i.e. density) and average degree (i.e. mean number of connections per node) of a network (Stam et al., 2014; van Wijk et al., 2010). Hence, comparing healthy and affected (or reorganized) brain networks might give biased results (Tewarie et al., 2015; van Wijk et al., 2010; Zalesky et al., 2010). A promising alternative analysis approach, which might solve these limitations, is selective assessment of the functional network backbone by means of the minimum spanning tree method. Network backbone analysis allows for unbiased comparison of networks since all minimum spanning trees have the same size and density (Stam et al., 2014; Tewarie et al., 2015). An increasing number of studies have shown the usefulness of this approach in capturing subtle network changes in brain development and ageing (Boersma et al., 2013; Otte et al., 2015; Smit et al., 2016; Vourkas et al., 2014) but also in brain diseases, like multiple sclerosis, Alzheimer’s disease and epilepsy (Engels et al., 2015; Tewarie et al., 2014; van Diessen et al., 2016, 2014).

In the present study we investigated the effects of prolonged periods of deafness on topological characteristics of brain functional network backbones. Therefore we acquired resting-state electroencephalograms from both healthy controls and congenital, pre-lingual and post-lingual deaf people in eyes-open and eyes-closed conditions. Opening and closing the eyes are very basic attention-directing behaviors (i.e. towards the internal or external world) (Xu et al., 2014), related to different brain states (Marx et al., 2004; Zhang et al., 2015). Moreover, deaf people completely lack audiovisual input during the eyes-closed condition. We made use of a unique homogeneous population in a representative rural region in sub-Saharan Africa where deafness is a common disability but cochlear implants are not available. The study of brain plasticity in deaf people in countries with a well-established health-care system is complicated as many people with hearing disabilities will have a cochlear implant. Cochlear implants changes the brain organization, including functional cortical reorganizations at rest (Strelnikov et al., 2010). If the implantation is done early, auditory language develops almost normally in those people (Hammes et al., 2002). In our study population none of the people had a cochlear implantation.

Electroencephalography recordings were used to construct functional networks and functional network backbones. We hypothesized enhanced functional connectivity between auditory and visual cortex in deaf people. In addition that functional network backbones differ between healthy and deaf people, indicating that sensory processing is a distributed brain function. As deaf people hardly get any sensory input when they are in resting-state condition with eyes closed, we speculated that we would detect larger network backbone-shifts between eyes-open and eyes-closed conditions in deaf people as compared to healthy controls. Lastly, we anticipated the backbone characteristics in deaf people to be related to years of American Sign Language (ASL) experience.

## Methods

### Study setting and ethics

The study pipeline as described below is visualized in **Figure 1**. Our study was conducted at two inclusive primary schools and one inclusive secondary school located at two separate rural places in Ebonyi State, southeast Nigeria in September-October 2016, which are part of a Community-Based Rehabilitation (CBR) program. In this inclusive education schools all community students, with or without disability, attend and are welcomed in age-appropriate, regular classes and are supported to learn, contribute and participate in all aspects of the life of the school. Standard ASL is taught at those Nigerian schools for more than twenty years. So all students, deaf and non-deaf learn sign language. This educational approach is a potential strategy to reduce the burden of disability (Eleweke and Rodda, 2002; Pförtner, 2014). The distribution of students with and without disabilities is about equal. The lessons are taught in English and if the teacher does not know sign language there is an interpreter for the deaf students. Every class consists of non-disabled as well as (a maximum of fifty percent) disabled students. Deaf children receive lessons in both Sign Language and Speech therapy. All children live in the surrounding villages for full integration within the community. In this way many people in the community master and/or understand Sign Language as well.

**Figure 1.**
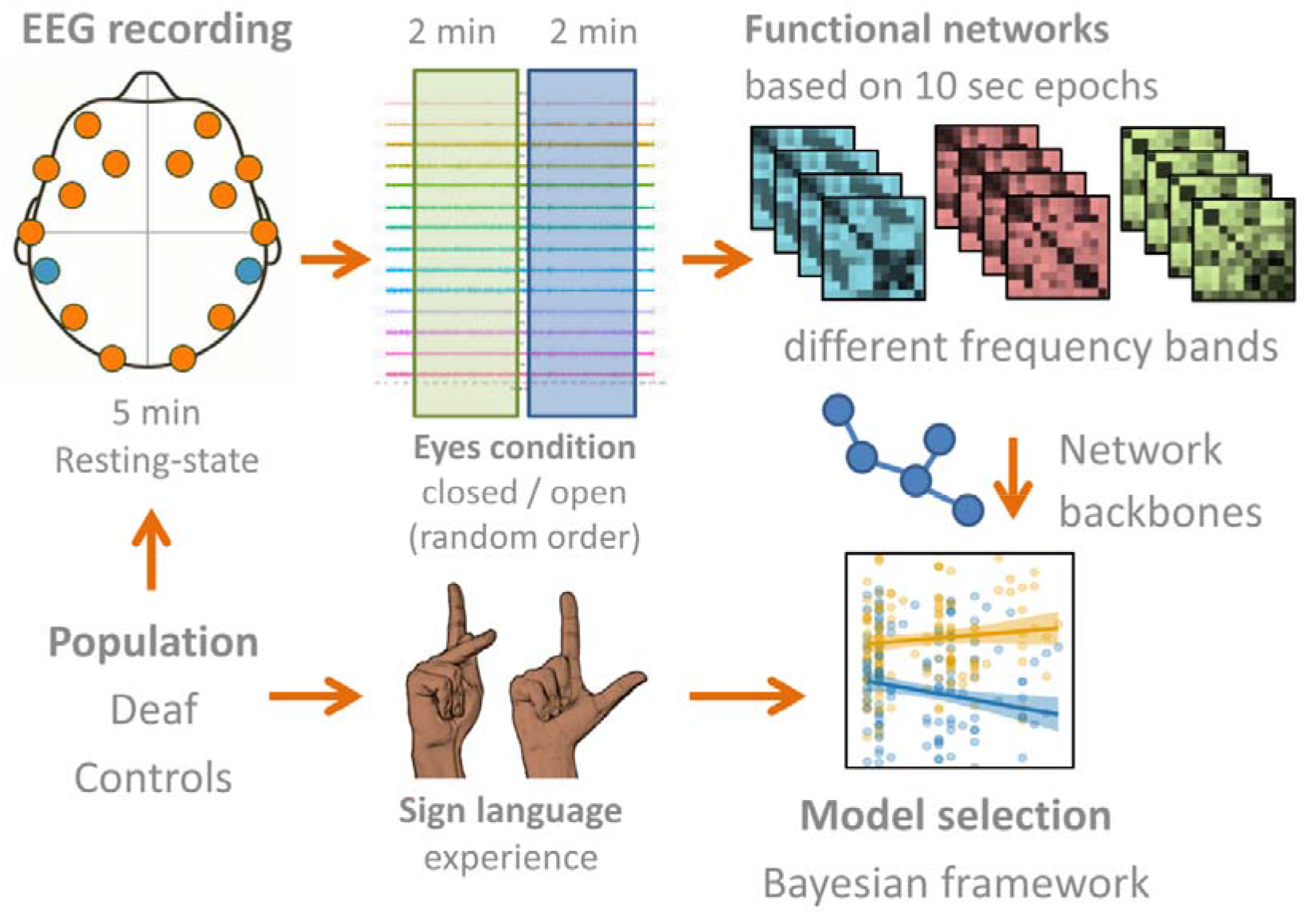
Schematic study protocol. Five minutes of resting-state electroencephalography (EEG) was acquired with wireless headsets in deaf and controls. The sensor locations corresponding to the fourteen channels are shown in orange; the CMS/DRL sensors are shown in blue. Two minutes within this recording were with eyes closed and two minutes with eyes open. The order of open/closed was random. Frequency bands included delta, theta, alpha, beta and gamma bands. Functional networks were constructed from 10-seconds epochs. Network backbone metrics were related with years of sign language experience. Interaction effects in the two populations and between ‘open/closed conditions’ were assessed with Bayesian model selection.

Our study was approved in by the organizational boards (RBC/CBR Effata) and the local health ministry (Izzi, Local Government Area) and federal government (Ebonyi State House of Assembly, Abakaliki [7-11-2016]) in Nigeria. Written informed consent was obtained from adult participants and caretakers of children below eighteen years. Assent was obtained from the children. Prior to asking volunteers for the recordings, the study protocol was clearly explained to all students in class.

### Participants

**Table 1** shows the demographic information of all participants. We included 193 participants between ten and 43 years old (mean age of 18.5 (standard deviation 6.0); gender: 103 male, 90 female), both students and teachers. Sign language experience of participants varied between zero and seventeen years at the time of recording. We selected two groups: hearing (n=122) and deaf (n=71) participants. The pre-lingual deaf participants were all capable of lip reading.

**Table 1.**
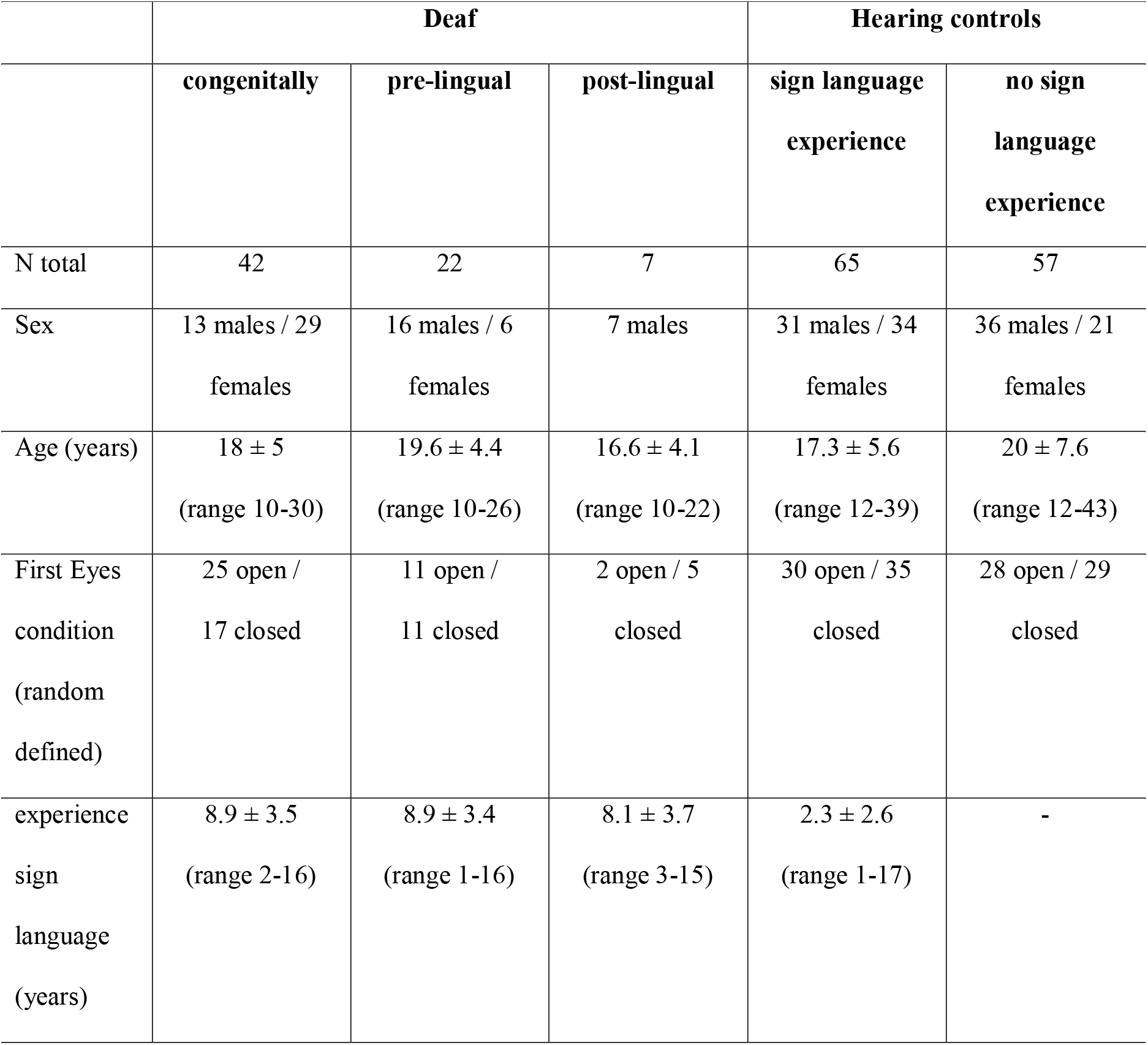
Participant characteristics.

### Data acquisition

We used a high-resolution sixteen sensor / fourteen channel EEG monitor configured to sample at 128 Hertz with a 16-bit resolution (EMOTIV Inc, San Francisco, USA) (Aspinall et al., 2013; Badcock et al., 2015; McMahan et al., 2015; Prause et al., 2016; Yu and Sim, 2016). This wireless headset can be connected to a computer via Bluetooth and is an invaluable tool to collect EEG signals from participants in rural or resource-limited areas, where access to a clinical type of EEG system is often impossible or burdensome. Two sensors were preserved for reference and grounding: the ‘common mode sense’ (CMS; located at P3) sensor was used as the active reference for absolute referencing. The ‘driven right leg’ (DRL; located at P4) sensor was used for feedback noise cancelation. The electrodes were located at anterofrontal (AF3, AF4, F3, F4, F7, F8), frontocentral (FC5, FC6), occipital (O1, O2), parietal (P7, P8) and temporal sites (T7, T8), according to the International 10–20 system. Signal quality scores are recorded for each electrode with a range of one to five (no units), with five as best quality.

Participants were asked to sit on a chair in a quiet room for five minutes while wearing the Emotiv headset. They were instructed to keep their eyes closed for the first three minutes and open in the next two minutes. The order of the conditions (i.e. ‘eyes closed’ or ‘eyes open’) was assigned alternatingly, so that half of the participants started in the ‘eyes open condition’ whereas the other half of the participants started in the ‘eyes closed condition’. The researcher kept a log on deviations from the protocol or unusual events in the environment that may affect the experiment. Example recordings are shown in **Figure 2**. The electroencephalography signals were band-pass filtered into the delta (0.5–4 Hz), theta (4–8 Hz), alpha (8–16 Hz), beta (16–32 Hz) and gamma (32–64 Hz) frequency bands (examples shown in Suppl. Figure 1 and 2).

**Figure 2.**
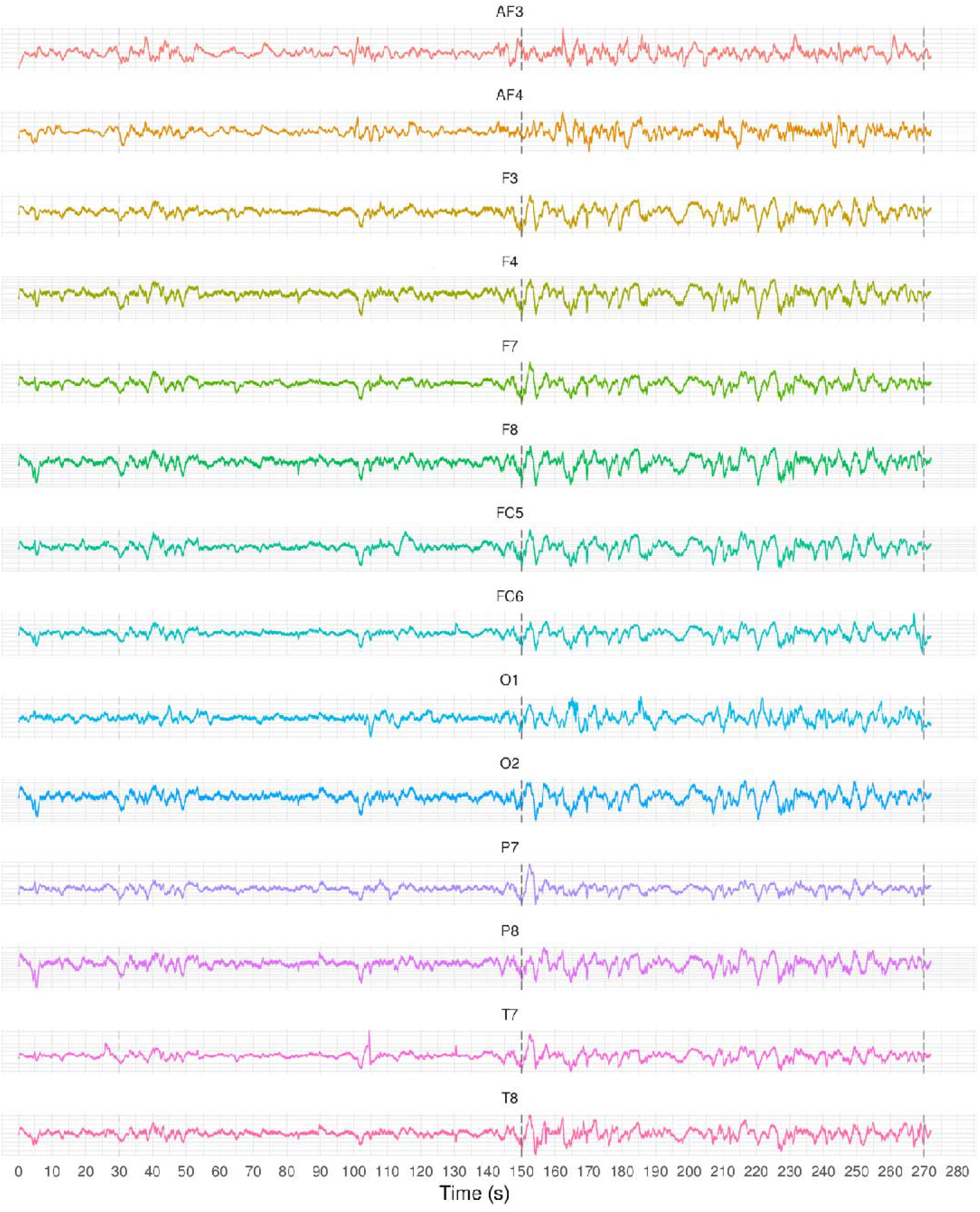
Raw EEG time-series from one participant (male, 22 years old, deaf). The y-scaling is arbitrary. Vertical grey lines indicate the initial acclimatization period, two minutes condition I (eyes closed in this subject) and two minutes condition II (eyes open). Labels of the fourteen channels are given above the time-series.

### Data cleaning and window selection

Time segments of the recordings were removed if i) the research log indicated a deviation from the protocol, ii) the electroencephalography signal quality score was below four for any of the channels, and iii) if the absolute deviation of the gyroscope signals relative to the gyroscope signal median was larger than five times the standard deviation. This threshold was based on visual data inspection (See example in **Suppl. Figure 3**). The cleaned filtered data were cut into 10-sec epochs. Functional connectivity and multiple network backbone metrics – calculated in a similar way as in our study – stabilize within recordings if the minimal epoch length is at least six seconds (Fraschini et al., 2016). Therefore we used this conservative 10-sec length. Multiple epochs per subject result in stable network backbone metrics (van Diessen et al., 2015).

### Functional connectivity

For each epoch a functional network was constructed. Recorded time-series within each epoch were used to determine functional connectivity between different electrodes capturing neuronal signals from underlying brain areas. Functional connectivity was computed and quantified with the phase lag index. This is a measure of the asymmetry of the distribution of instantaneous phase differences between two time series and scales between zero and one (Pillai and Sperling, 2006). It is relative resistant to the influence of common sources, including volume conduction and active reference electrodes. An index of zero indicates no phase coupling between time series, or coupling with a phase difference centered on zero ± p radians. A non-zero index indicates the presence of phase coupling. A more mathematical description of computing the phase lag index can be found elsewhere (Stam et al., 2007).

### Occipital – parietal functional connectivity

We expected remodeling of the auditory cortex in deaf people reflected as enhanced synchronization between the auditory and visual cortex. We therefore characterized in both groups the average phase lag index between the electrodes P7/P8, covering the auditory cortex, and the occipital electrodes O1/O2, covering the visual cortex, at eyes-open and eyes-closed conditions.

### Minimum spanning tree analysis

For each functional network a minimum spanning tree (MST) was calculated from the connectivity graph *G* by applying Kruskal’s algorithm (Kruskal, 1956). An MST captures the network’s backbone and is defined as a subset of the network nodes (forming the original weighted graph *G*) that connects all the nodes and does not contain cycles or loops (Jackson and Read, 2010). A minimum spanning tree *T* minimizes the sum of the costs of its edges, 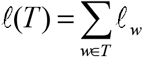, over the set of all possible spanning trees on *G* (Hidalgo et al., 2007).

The following MST metrics were calculated at nodal or network level:

i. *Maximum node degree (nodal):* We summarized every tree by taking the maximum node degree: S_max_. Which is the node with the maximum number of connections.
ii. *Leaf number (N_leaf_) (network):* the number of nodes of the tree with exactly one connection to any other node (with maximum degree = 1). A higher leaf number is related to increased global efficiency and integration (Stam et al., 2014; Tewarie et al., 2015).
iii. *Diameter (d) (network):* the largest distance between any two nodes in a tree, which has a lower bound of two and an upper bound of m = N – 1. The largest possible diameter will decrease with increasing leaf number (Boersma et al., 2013; Stam et al., 2014; Tewarie et al., 2015).
iv. *Eccentricity (network):* the shortest path length between a tree node *I* and any other node from the tree. Eccentricity decreases when nodes become more central in the tree.
v. *Radius (nodal):* the smallest node eccentricity in the tree. The lower the eccentricity, the more central a node in a tree.
vi. *Strength (nodal):* the tree node strength is a summation of all nodal connection weights (Hagmann et al., 2010; Rubinov and Sporns, 2010).
vii. *Maximum betweenness centrality (BC_max_)*: a network hub metric which relies on the identification of the number of shortest paths that pass through a node (Rubinov and Sporns, 2010). The more the passages, the higher the betweenness-centrality (i.e. hubness), which is defined by 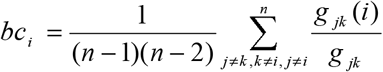 where *g_jk_* is the shortest path between two nodes and *g_jk_(i)* is the number of node paths that actually pass through *i*. We summarized the tree by taking the maximum betweenness centrality: *BC_max_*.
viii. Closeness centrality (nodal): the inverse of the sum of all distances to other nodes (Sabidussi, 1966).

### Statistical analyses

We compared topological network differences between congenitally deaf, pre- and post lingually deaf and hearing subjects by means of Bayes factors extracted from Bayesian model comparisons (**Table 2**). We determined the model likelihood of a null model without interaction between group and eye condition and the likelihood of a model with an interaction between group and eye condition for each MST network metric. Bayes factors give the ratio of model likelihoods; thereby providing which model (i.e. the presence or absence of a difference in the parameter change) is supported (i.e. more likely to occur). This was repeated for each frequency band separately. Since sex (Boersma et al., 2011) and age (Smit et al., 2012) influence functional network topologies, we included both as covariates in the analysis to correct for potential sex- and age-related group effects.

**Table 2.** Bayes factors and their interpretations, based on (Raftery, 1995). M_0_: the baseline model without interaction term, M_1_: the alternative model with interaction term. In the subsequent tables the Bayes factors for M_1_ are reported only.

| log (Bayes factor) |  |  | Interpretation |
| --- | --- | --- | --- |
| | > | 100 | Extreme evidence for $M_1$ |
| 30 | – | 100 | Very strong evidence for $M_1$ |
| 10 | – | 30 | Strong evidence for $M_1$ |
| 3 | – | 10 | Moderate evidence for $M_1$ |
| 1 | – | 3 | Anecdotal evidence for $M_1$ |
|  | 1 |  | No evidence |
| 1/3 | – | 1 | Anecdotal evidence for $M_0$ |
| 1/10 | – | 1/3 | Moderate evidence for $M_0$ |
| 1/30 | – | 1/10 | Strong evidence for $M_0$ |
| 1/100 | – | 1/30 | Very strong evidence for $M_0$ |
| | < | 1/100 | Extreme evidence for $M_0$ |

All network analyses, statistical modeling and visualization were performed in R (http://www.r-project.org/) using the packages *igraph, BayesFactor* and *ggplot2.* All epoch data is available at the Open Science Framework in anonymized form (Otte et al., 2018).

## Results

### Occipital – parietal functional connectivity

Functional connectivity differences between eyes-open and eyes-closed in occipital-parietal cortex are shown in **Figure 3: left panel**. The functional connectivity is lower in the eyes-open condition in the theta, alpha and beta frequency range. Differences are most pronounced in the alpha frequency range with −54.9% (95% confidence interval (CI): −68.2 to −41.6%) reduction in functional connectivity during switching from eyes-closed to eyes-open in the control group (**Figure 3: right panel**). This reduction was larger in the deaf group −88.0% (CI: −112.4 to −63.6%). Reductions in functional connectivity between eyes-closed and eyes-open were also present in the beta frequency range: −27.7% (CI: −37.3 to −18.0) in controls and −36.1% (CI: −49.2 to −23.0) in deaf people. These alpha and beta functional connectivity reductions were statistically significant (**Table 3**).

**Table 3.** Bayes factors of the interaction between condition (i.e. eyes-closed and eyes-open) and the group (i.e. deaf and controls) per frequency band.

| *Frequency* | **Bayes factor** |
| --- | --- |
| Delta (0.5-4 Hz) | 0.04 |
| Theta (4-8 Hz) | 0.07 |
| Alpha (8-16 Hz) | **922.92** <sup>v</sup> |
| Beta (16-32 Hz) | **2.42** <sup>I</sup> |
| Gamma (32-64 Hz) | 0.03 |

<sup>I</sup> *Anecdotal evidence*, <sup>v</sup> *Extreme evidence for interaction*

**Figure 3.**
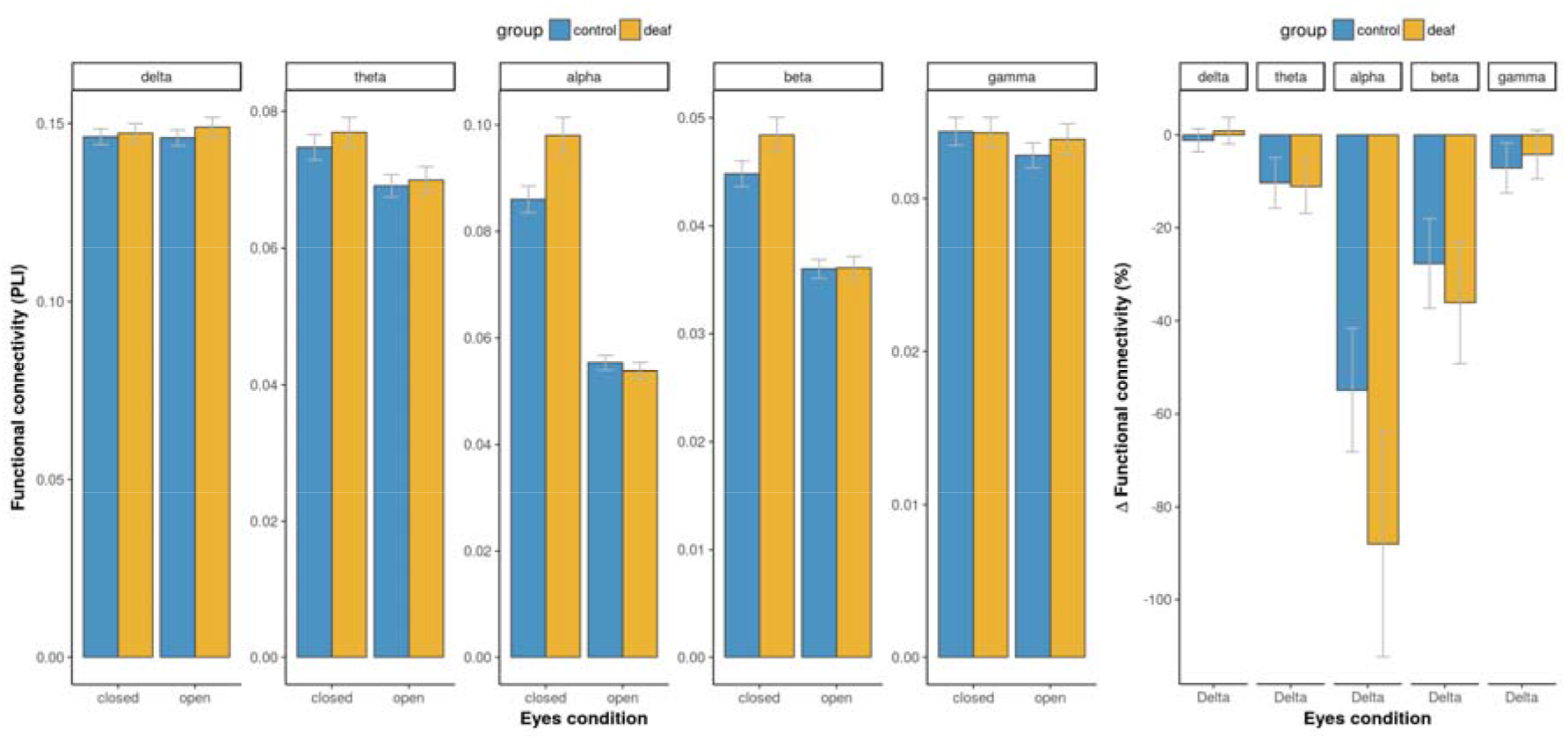
Functional connectivity between the occipital and parietal cortex. *Left panel:* The average functional connectivity, quantified with the phase lag index, between O1/O2 and P7/P8 (y-axis) is shown for eyes-open and eyes-closed conditions (x-axis) and all frequency bands (top) in controls and deaf people. *Right panel:* The delta functional connectivity between eyes-open en eyes-closed is plotted as percentage change relative to the eyes-open functional connectivity values for each frequency band, based on the data shown in the left panel: Δ in % = 100 × (closed – open) / open. Error bars represent the 95% confidence intervals.

### Functional backbone differences between eyes-open and eyes-closed conditions

Transition from eyes-closed condition to eyes-open condition showed visible changes in functional network backbone characteristics. The most notable differences were found in the alpha band and beta band, which are shown in **Figure 4**. In both the alpha and beta band the backbone leaf number and kappa were lower in eyes-open condition than in eyes-closed condition for both deaf and controls. Contrary, in both the alpha and beta band the backbone diameter, eccentricity and radius were higher in eyes-open condition compared to eyes-closed condition, again for both deaf and controls. Furthermore, changing from eyes-closed to eyes-open condition showed an increase in closeness centrality and a decrease in mean and maximum node strength, in the alpha band only.

**Figure 4.**
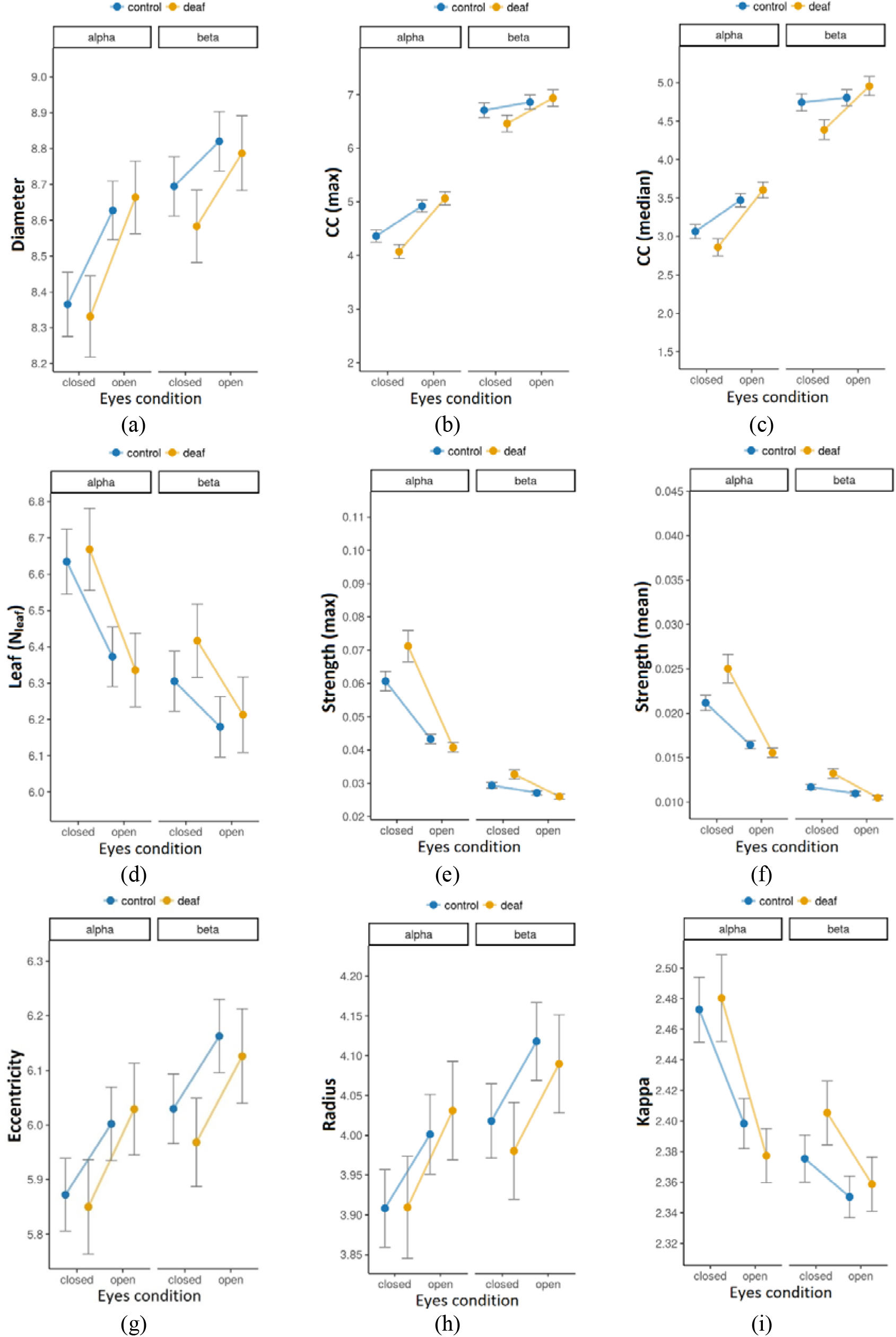
Network backbone comparisons between eyes open and eyes closed conditions for both deaf and controls. Functional network backbone characteristics in the alpha band (8-16Hz) and beta band (16-32 Hz) (top), are shown for deaf (yellow) and controls (blue) for both the eyes open and eyes closed condition (x-axis) and indicated by the following minimum spanning tree metrics (y-axis): (a) diameter, (b) maximum closeness centrality, (c) median closeness centrality, (d) leaf number, (e) maximum strength, (f) median strength, (g) eccentricity, (h) radius and (i) kappa. Error bars represent the 95% confidence intervals.

### Larger functional network modifications in deaf

As shown in **Figure 4**, from the eyes-open to the eyes closed condition, several functional network backbone characteristics show larger shifts in deaf as compared to hearing participants. As indicated by the Bayes factors in **Table 4**, there is extreme evidence for a larger shift (i.e. from eyes open to eyes closed condition) of the functional backbone node strength in both alpha and beta band in deaf. Furthermore in the alpha band there was moderate to strong evidence for a larger shift of the backbone closeness centrality median and maximum respectively in deaf. In addition, in the beta band there was some evidence for a larger shift in backbone closeness centrality, leaf number, diameter and kappa in deaf as compared to controls.

**Table 4.** Bayes factors of the interaction between conditions (i.e. eyes-closed and eyes-open) and the group (i.e. deaf and controls) per frequency band.

|  | *Delta band* | *Theta band* | *Alpha band* | *Beta band* | *Gamma band* |
| --- | --- | --- | --- | --- | --- |
| **MST-metric** | **(0.5-4 Hz)** | **(4-8 Hz)** | **(8-16 Hz)** | **(16-32 Hz)** | **(32-64 Hz)** |
| Strength (max) | 0.06 | 0.08 | **1547.06<sup>V</sup>** | **8364.70<sup>V</sup>** | 0.13 |
| Strength (mean) | 0.11 | 0.07 | **24539.81<sup>V</sup>** | **224.19<sup>V</sup>** | 0.15 |
| Degree (max) | 0.03 | 0.08 | 0.05 | **4.64<sup>II</sup>** | 0.12 |
| BC (max) | 0.02 | 0.06 | 0.06 | 0.32 | 0.06 |
| BC (median) | 0.05 | 0.08 | 0.05 | 0.23 | 0.10 |
| CC (max) | 0.04 | 0.07 | **26.72<sup>III</sup>** | 0.64 | 0.09 |
| CC (median) | 0.03 | 0.08 | **7.17<sup>II</sup>** | **1.13<sup>I</sup>** | 0.09 |
| Leaf | 0.05 | 0.06 | 0.06 | **1.11<sup>I</sup>** | 0.07 |
| Diameter | 0.05 | 0.06 | 0.06 | **1.10<sup>I</sup>** | 0.07 |
| Eccentricity | 0.04 | 0.05 | 0.05 | 0.40 | 0.05 |
| Radius | 0.05 | 0.06 | 0.06 | 0.23 | 0.06 |
| Tree-hierarchy | 0.04 | 0.06 | 0.06 | 0.27 | 0.07 |
| Kappa | 0.04 | 0.10 | 0.06 | **2.63<sup>I</sup>** | 0.07 |

*BC = betweenness centrality, CC = closeness centrality, <sup>I</sup> Anecdotal evidence, <sup>II</sup> Moderate evidence, <sup>III</sup> Strong evidence, <sup>V</sup> Extreme evidence for interaction*

Furthermore, the results showed differences among the deaf participants. **Figure 5** shows a difference in functional backbone node strength between different types of deafness (i.e. congenital, pre-lingual and post-lingual deafness), which is supported by the Bayes factor indicating strong evidence as shown in **Table 5**. In addition, these Bayes factors show moderate evidence for a difference among deaf participants in both the maximum and the median of closeness centrality of functional network backbones.

**Table 5.**
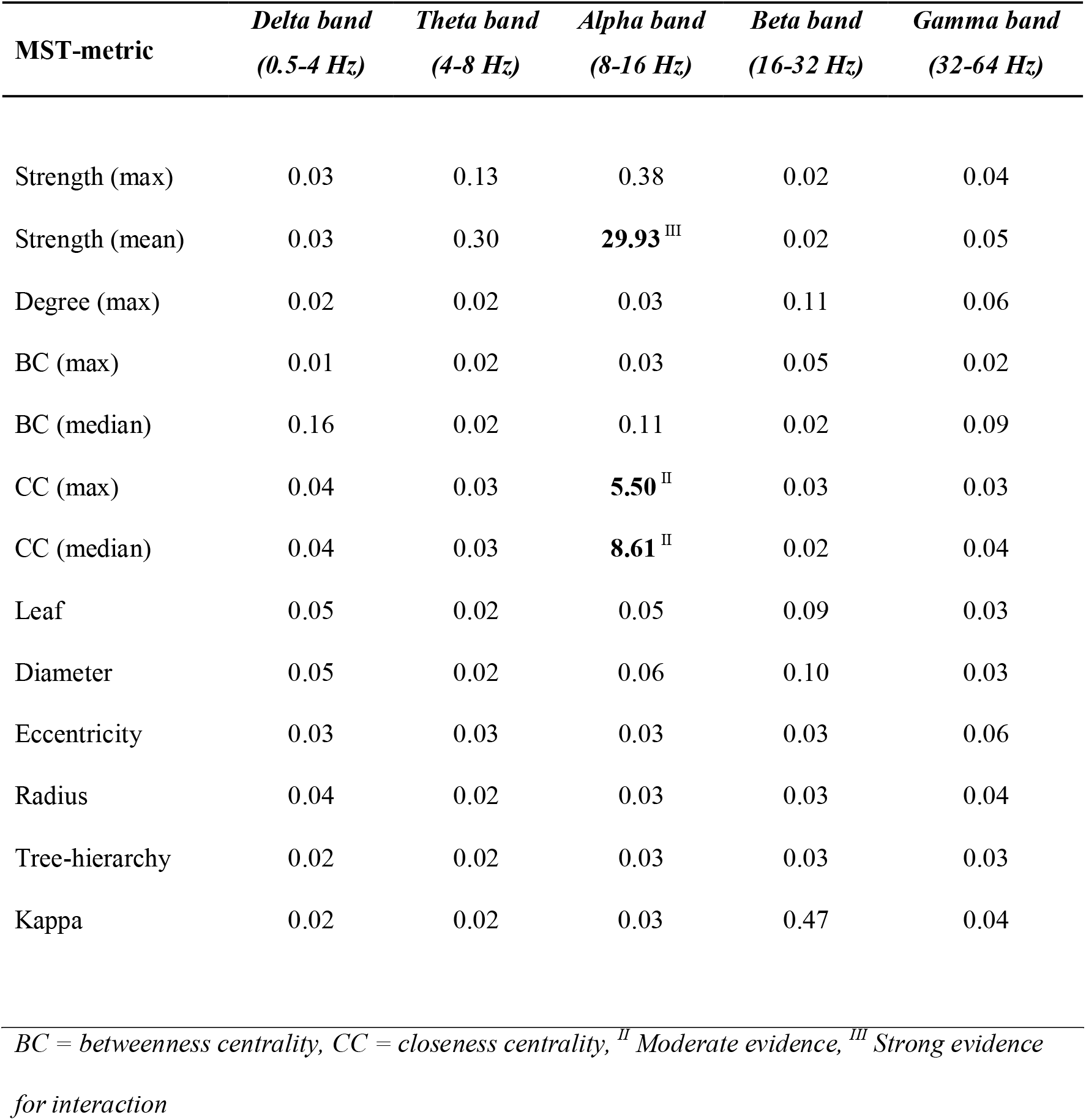
Bayes factors of the interaction between condition (i.e. eyes-open and eyes-closed) and type of deafness (i.e. congenital, pre-lingual and post-lingual) per frequency band.

| **MST-metric** | *Delta band*<br>*(0.5-4 Hz)* | *Theta band*<br>*(4-8 Hz)* | *Alpha band*<br>*(8-16 Hz)* | *Beta band*<br>*(16-32 Hz)* | *Gamma band*<br>*(32-64 Hz)* |
| --- | --- | --- | --- | --- | --- |
| Strength (max) | 0.03 | 0.13 | 0.38 | 0.02 | 0.04 |
| Strength (mean) | 0.03 | 0.30 | **29.93<sup>III</sup>** | 0.02 | 0.05 |
| Degree (max) | 0.02 | 0.02 | 0.03 | 0.11 | 0.06 |
| BC (max) | 0.01 | 0.02 | 0.03 | 0.05 | 0.02 |
| BC (median) | 0.16 | 0.02 | 0.11 | 0.02 | 0.09 |
| CC (max) | 0.04 | 0.03 | **5.50<sup>II</sup>** | 0.03 | 0.03 |
| CC (median) | 0.04 | 0.03 | **8.61<sup>II</sup>** | 0.02 | 0.04 |
| Leaf | 0.05 | 0.02 | 0.05 | 0.09 | 0.03 |
| Diameter | 0.05 | 0.02 | 0.06 | 0.10 | 0.03 |
| Eccentricity | 0.03 | 0.03 | 0.03 | 0.03 | 0.06 |
| Radius | 0.04 | 0.02 | 0.03 | 0.03 | 0.04 |
| Tree-hierarchy | 0.02 | 0.02 | 0.03 | 0.03 | 0.03 |
| Kappa | 0.02 | 0.02 | 0.03 | 0.47 | 0.04 |

*BC = betweenness centrality, CC = closeness centrality, <sup>II</sup> Moderate evidence, <sup>III</sup> Strong evidence for interaction*

**Figure 5.**
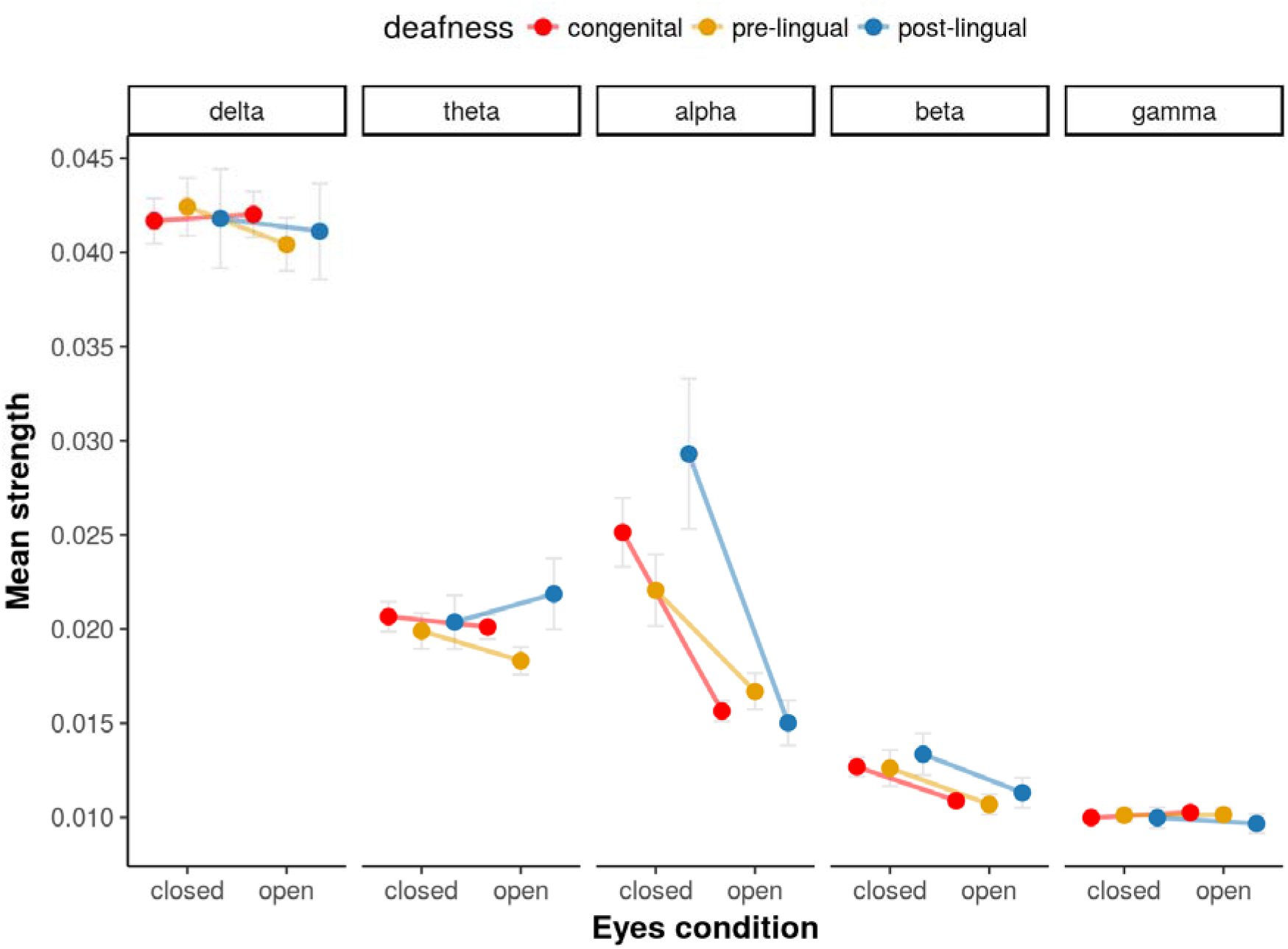
Average functional backbone node strength differs across types of deafness. The average functional backbone node strength (y-axis) shown for eyes-open and eyes-closed conditions (x-axis) and all frequency bands (top) differed across different types of deafness, i.e. congenital (red), pre-lingual (yellow) and post-lingual (blue). Error bars represent the 95% confidence intervals.

### Relation between backbone characteristics and American Sign Language

Next we investigated the effect of ASL on functional backbone characteristics, which revealed a correlation between experience in ASL and several functional backbone metrics. Initially, no distinction was made between deaf and hearing subjects, because hearing subjects had relatively little ASL experience. **Figure 6** shows that ASL experience is related to altered backbone characteristics in the theta band. More specifically, an increase in ASL experience is related to a larger backbone diameter and radius, a higher backbone eccentricity and betweenness centrality (median), combined with a lower leaf number, kappa and hierarchy (see **Table 6** for corresponding Bayes factors). **Figure 7** shows the relationship between ASL experience and functional backbone characteristics for deaf subjects only. An increase in ASL experience is related to a larger maximal closeness centrality in the theta-band network (see **Table 7** for corresponding Bayes factors). Furthermore, **Table 6** shows strong evidence for a correlation between mean and maximum backbone strength and ASL experience as well as extreme evidence for a relation between closeness centrality and ASL experience in the delta band.

**Table 6.** Bayes factors of the relation between the MST-metric and ASL-experience.

|  | *Delta band* | *Theta band* | *Alpha band* | *Beta band* | *Gamma band* |
| --- | --- | --- | --- | --- | --- |
| **MST-metric** | **(0.5-4 Hz)** | **(4-8 Hz)** | **(8-16 Hz)** | **(16-32 Hz)** | **(32-64 Hz)** |
| Strength (max) | **31.37<sup>IV</sup>** | 0.05 | **1.14<sup>I</sup>** | 0.09 | 0.05 |
| Strength (mean) | **26.10<sup>III</sup>** | 0.05 | **1.55<sup>I</sup>** | 0.06 | 0.07 |
| Degree (max) | 0.04 | **2.80<sup>I</sup>** | 0.08 | 0.10 | 0.06 |
| BC (max) | 0.03 | 0.46 | 0.05 | 0.04 | 0.04 |
| BC (median) | 0.07 | **276.98<sup>V</sup>** | 0.04 | 0.05 | 0.04 |
| CC (max) | **106.82<sup>V</sup>** | 0.41 | 0.09 | 0.04 | 0.10 |
| CC (median) | **196.19<sup>V</sup>** | 0.24 | 0.10 | 0.04 | 0.08 |
| Leaf | 0.15 | **1.450.69<sup>V</sup>** | 0.04 | 0.14 | 0.05 |
| Diameter | 0.16 | **1.474.65<sup>V</sup>** | 0.04 | 0.14 | 0.05 |
| Eccentricity | 0.04 | **70.93<sup>IV</sup>** | 0.07 | 0.04 | 0.05 |
| Radius | 0.05 | **33.39<sup>IV</sup>** | 0.05 | 0.05 | 0.07 |
| Tree-hierarchy | 0.51 | **14.90<sup>III</sup>** | 0.05 | 0.14 | 0.05 |
| Kappa | 0.06 | **225.97<sup>V</sup>** | 0.04 | 0.38 | 0.05 |

*BC = betweenness centrality, CC = closeness centrality, <sup>I</sup> Anecdotal evidence, <sup>III</sup> Strong evidence,*
*<sup>IV</sup> Strong evidence, <sup>V</sup> Extreme evidence for relation*

**Table 7.** Bayes factors of the relation between the MST-metric and ASL-experience in deaf people only.

| **MST-metric** | ***Delta band*** | ***Theta band*** | ***Alpha band*** | ***Beta band*** | ***Gamma band*** |
| --- | --- | --- | --- | --- | --- |
|  | ***(0.5-4 Hz)*** | ***(4-8 Hz)*** | ***(8-16 Hz)*** | ***(16-32 Hz)*** | ***(32-64 Hz)*** |
| Strength (max) | 0.26 | 0.14 | 0.07 | 0.07 | 0.08 |
| Strength (mean) | 0.48 | 0.33 | 0.06 | 0.08 | 0.07 |
| Degree (max) | 0.03 | 0.14 | 0.08 | 0.06 | 0.39 |
| BC (max) | 0.08 | 0.97 | 0.06 | 0,10 | 0.07 |
| BC (median) | 0.13 | 0.12 | 0.12 | 0.06 | 0.13 |
| CC (max) | **1.53**<sup>I</sup> | **1.61**<sup>I</sup> | 0,07 | 0.06 | 0.06 |
| CC (median) | **1.64**<sup>I</sup> | 0.92 | 0.08 | 0.05 | 0.06 |
| Leaf | 0.15 | 0.13 | 0.20 | 0.07 | 0.15 |
| Diameter | 0.16 | 0.13 | 0.20 | 0,07 | 0.16 |
| Eccentricity | 0.04 | 0.12 | 0.09 | 0.06 | 0.07 |
| Radius | 0.05 | 0.09 | 0,06 | 0.06 | 0.08 |
| Tree-hierarchy | 0.7 | 0.06 | 0.19 | 0,12 | 0.13 |
| Kappa | 0.04 | 0.15 | 0.24 | 0.06 | 0.25 |

*BC = betweenness centrality, CC = closeness centrality, <sup>I</sup> Anecdotal evidence for relation*

**Figure 6.**
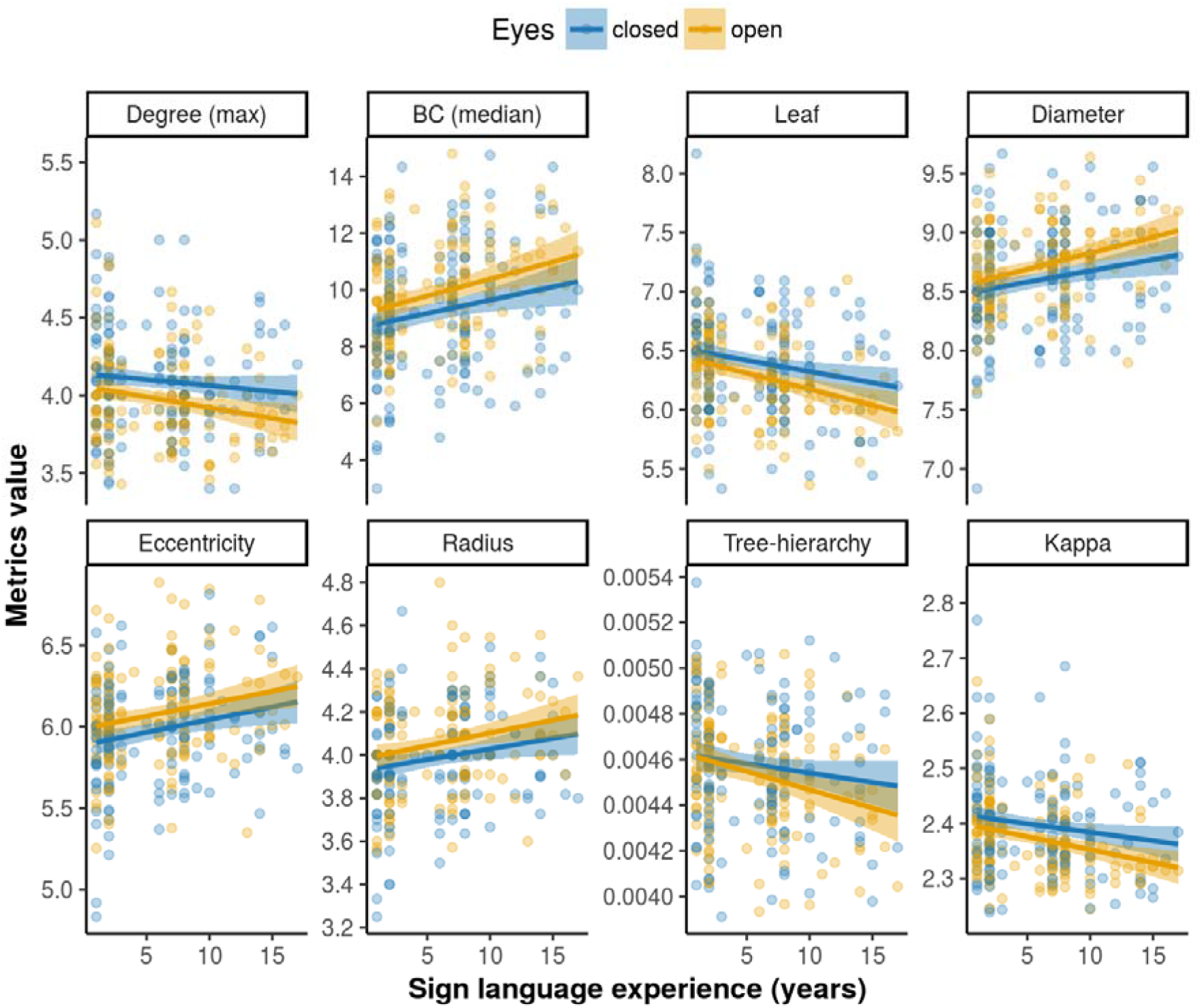
The relation between American Sign Language (ASL) and functional backbone characteristics. The ASL experience in years (x-axis) related to functional backbone characteristics (y-axis) in the theta band (4-8 Hz) for both the eyes open (yellow) and eyes closed (blue) condition, as indicated by the following minimum spanning tree metrics (from top-left to bottom-right): maximum degree, median betweenness centrality (BC), leaf number, diameter, eccentricity, radius, tree-hierarchy, and kappa. Shaded areas: 95% confidence intervals.

**Figure 7.**
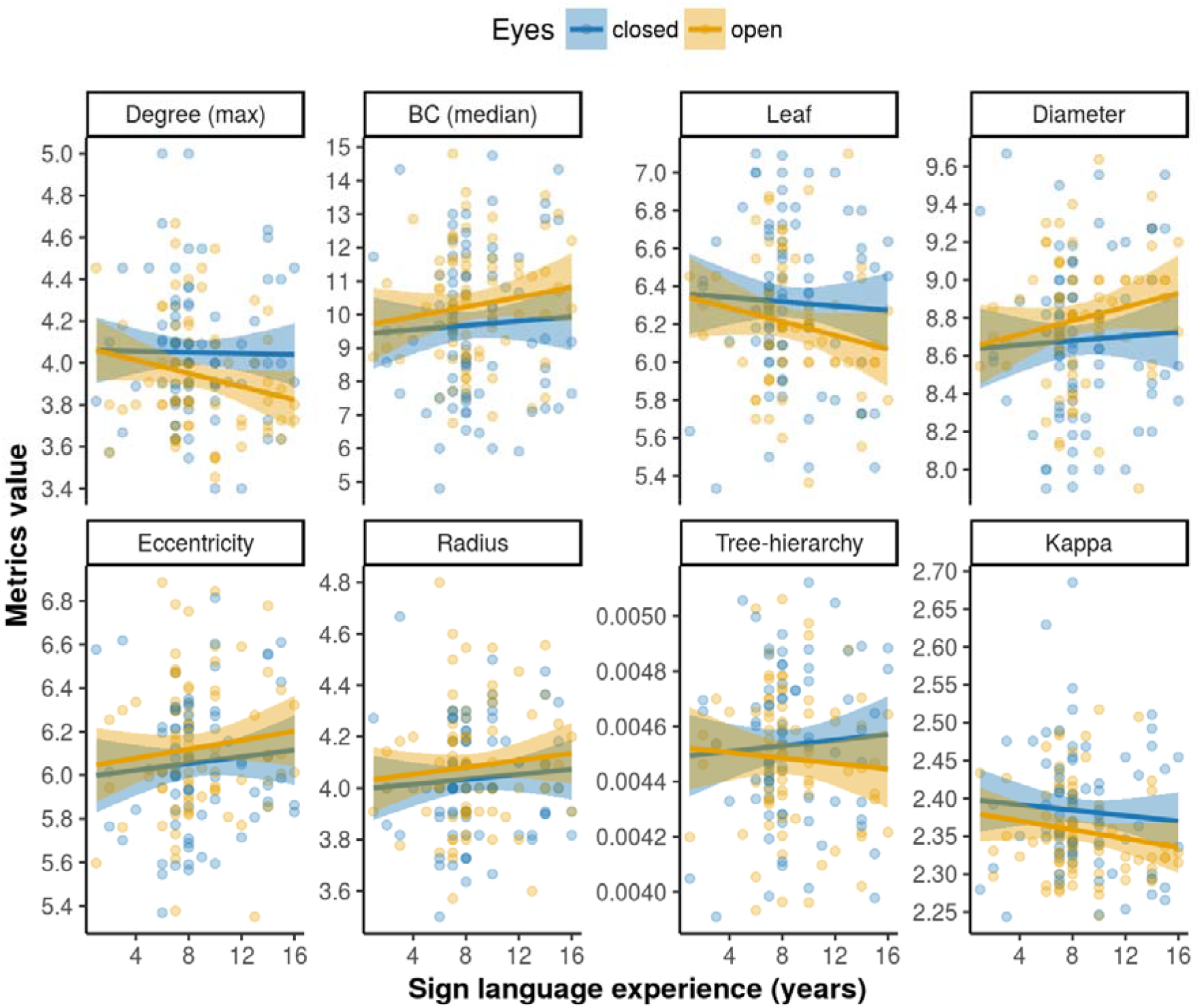
The relation between American Sign Language (ASL) and functional backbone characteristics in deaf subject only. The ASL experience in years (x-axis) related to functional backbone characteristics (y-axis) in the theta band (4-8 Hz) for both the eyes open (yellow) and eyes closed (blue) condition, as indicated by the following minimum spanning tree metrics (from top-left to bottom-right): maximum degree, median betweenness centrality (BC), leaf number, diameter, eccentricity, radius, tree-hierarchy, and kappa. Shaded areas: 95% confidence intervals.

## Discussion

Our study employed resting-state EEG to investigate and map functional network backbone differences between deaf and hearing people. We showed that transition from the eyes-closed to the eyes-open condition was associated with changes in functional connectivity between occipital and parietal cortex as well as changes in functional network backbones. These occur especially in the alpha and beta frequency bands, in both deaf and hearing participants. This indicates increased synchronization between auditory and visual cortical regions in response to visual stimuli (Boytsova and Danko, 2010) or suggests topological network reorganization.

### Functional cortical remapping

The ‘spontaneous’ intrinsic activity of the brain is affected by incoming stimuli, such as visual information, which will be different in the ‘eyes open’ and ‘eyes closed’ conditions. Moreover, the number and strength of connections also change, which is known as ‘alpha synchronization’ and is shown to be related to the wiring cost variation between ‘eyes-open’ en ‘eyes-closed’ conditions (Gómez-Ramírez et al., 2017). However, in addition to connectivity variation between conditions, the strong interaction effect we found between deaf and control conditions in alpha and beta functional connectivity between occipital and parietal cortex favors the hypothesis that in grown-up humans the neocortex possesses longterm plasticity. This cortical plasticity is known to be a multicomponent process which involves both synaptic and cellular mechanisms (Feldman, 2009). What is interesting from our results is that the modifications in cortical mapping occur at a relative large distance, namely between occipital and parietal regions. This remodeling is also in line with neural plasticity of the primary visual cortex. It has been shown in multiple studies across three different species, such as cats and monkeys, that the location of V1 neuronal receptive fields shifts after induction of lesions in the retina (Calford et al., 2000; Chino et al., 1995; Heinen and Skavenski, 1991; Kaas et al., 1990). These shifts are cortically mediated given that the thalamic input to the visual cortex shows limited reorganization (Eysel, 1982). In deaf people it may even be the case, as suggested by our increased neuronal synchronization between occipital and parietal recordings, that these shifts occur at a much larger scale as well. Validation studies with experimental animal models are required to characterize the underpinnings of this enhanced functional synchronization.

### Functional backbone differences between eyes-open and eyes-closed conditions

During the eyes-open condition the functional network backbone showed a larger diameter, higher eccentricity and a lower leaf number in the alpha and beta bands. This suggests that the backbone topology in the eyes-open condition was more chainlike (i.e. less integration and global efficiency), whereas during the eyes-closed condition the topology was more star-like (i.e. more functional integration and global efficiency) (Stam et al., 2014). Visual input during eyes-open conditions might suppress the connectivity of default-mode network activity during eyes-closed condition (Chen et al., 2008). These results may also be explained by alpha desynchronization, which will lead to less connections in general, indicating a ‘real change’ in functional connectivity and not just a change in alpha activity (Gómez-Ramírez et al., 2017). The alpha desynchronization from the eyes-open to the eyes closed condition in deaf as well as healthy subjects is in line with previous research among healthy adults (Barry et al., 2007; Gómez-Ramírez et al., 2017) and children (Barry et al., 2009). Two other studies reported increased global efficiency of functional networks in the alpha band, but not in the beta band, during the eyes-open condition as compared to the eyes-closed condition (Miraglia et al., 2016; Tan et al., 2013), which seem to contradict our findings. These discrepancies might be explained by demographical and methodological differences. Miraglia et al. (2016) only included 30 healthy elderly with an average age of 65.4, while Tan and colleagues included only 21 healthy Chinese university students, whereas our study includes 193 participants, both teenagers and adults. Both studies used more EEG electrodes or regions of interest (i.e. respectively 128 and 84) leading to larger networks with potential higher densities. Moreover, these previous studies used classical graph analysis to investigate whole brain network, whereas we have used MST analysis and studied the functional backbones of neural networks only, which might arguably lead to slightly different results.

### Larger functional network modifications in deaf

We found larger functional backbone shifts in both alpha and beta bands, from the eyes-open to the eyes-closed condition in deaf people compared to hearing controls. These differences between the eyes-open and eyes-closed conditions may be related to the different attentional states of the brain (i.e. internal versus external attention) (Marx et al., 2004; Xu et al., 2014; Zhang et al., 2015). In the beta band, the shift (i.e. from a more chainlike backbone in the eyes-open condition towards a more star-like backbone in eyes-closed condition) was more pronounced in deaf than in healthy controls. Specifically we found a larger shift of functional backbone node strength in deaf people from eyes-open to eyes-closed, indicating a higher overall physiological efficacy of backbone nodes in deaf during the eyes closed condition, but a lower overall efficacy during the eyes open condition (Hagmann et al., 2010). These effects might be more pronounced in deaf due to cross-modal plasticity (Bavelier et al., 2006; Bavelier and Neville, 2002; Hauser et al., 2007; Merabet and Pascual-Leone, 2010), especially in the alpha band, which has been related to visual stimuli (Boytsova and Danko, 2010) and awareness (Putilov and Donskaya, 2014), as well as the complete lack of audiovisual input in the eyes-closed condition.

The larger functional backbone shifts we found in deaf people, especially in the alpha band, may therefore reflect increased awareness in eyes-open condition (i.e. they rely to a greater extend on visual modality) and decreased awareness in eyes-closed condition. This hypothesis is based on the deprived sensory input typically found in deaf people. Interestingly, we also found that backbone node strength differed across the different forms of deafness and might depend on whether people were born deaf (i.e. congenital deafness) or acquired deafness later in life (i.e. pre-lingual or post-lingual deafness).

The lack of differences between deaf and hearing in functional backbone shifts from the eyes-open to the eyes closed condition in other frequency bands (i.e. delta, theta and gamma), suggests that auditory deprivation does not alter functional networks as much in these frequency bands. This might indicate that auditory deprivation leads to a strengthened functional networks regarding situations related to sensory input (i.e. the alpha and beta bands), rather than functional networks consisting of gamma waves. Gamma rhythms are involved in higher processing tasks and cognitive functioning (Herrmann et al., 2010)), whereas theta waves are involved in daydreaming, sleep and creativity. It seems therefore that topology of functional networks in the brain (of deaf) is strongly related to conscious achievement together with visual input, although further research is needed to investigate this into more detail.

### Relation between backbone characteristics and American Sign Language

Another interesting finding of our study is the clear relation between years of ASL experience and functional backbone characteristics in the theta band. Increasing experience in sign language reduced the leaf number, the maximum degree and the hierarchy of the functional backbones, while it increased the eccentricity, diameter, median betweenness centrality and radius of functional backbones (i.e. together indicating less integration and efficiency). This means that the organization of functional backbones shifts toward a more chain-like or decentralized organization with gaining ASL experience. This pattern is unlikely to be caused by age alone as these functional network backbone patterns have different or even opposing (e.g. leaf number, tree-hierarchy and diameter) directions in studies across the lifespan (Smit et al., 2016). The correlation between functional backbone characteristics and years of ASL experience is in line with previous research showing plasticity of functional and structural network organization in the brain due to sign language (Meyer et al., 2007). However, to the best of our knowledge there is no study that includes early-deaf teenagers and adults with sign language experience ranging from one year up to sixteen years.

### Advantages of study design and tools

Our results again show that MST graph parameters are highly suitable in exploring the topology and connectivity of brain networks (Engels et al., 2015; Tewarie et al., 2014; van Diessen et al., 2016), in our case related to cross modal neuroplasticity. Also our study shows the usefulness of the portable electroencephalography device as an invaluable tool to be used in rural or resource-limited African areas. This device enabled us to acquire an unique data-set of recordings from deaf and hearing subjects in a country where auditory deprivation is more prevalent than in Western countries, but where neuroimaging research is often impossible or burdensome due to geographic conditions, poor health-care infrastructure and high levels of poverty.

### Study limitations and future directions

The downside of electroencephalography is that neural activity is measured at a few places on the scalp and lacks information from deeper brain structures. In addition, the current data set was limited to fourteen channels. The signals are linear combinations of the neural generators they project to the scalp location of the electrodes. Together this means a reduction in precision of functional network mapping. Other neuroimaging techniques, such as MRI, might therefore be more capable to capture different activation patterns of the whole brain, specifically in sub-cortical areas, also because functional MRI studies have shown that visual motion stimuli caused a response in some specific auditory regions in deaf subjects (Almeida et al., 2015; Ding et al., 2015; Karns et al., 2012; Li et al., 2015; Neville et al., 1998; Petitto, 2000; Sadato et al., 2005, 2004; Scott et al., 2014; Shiell et al., 2016; Smith et al., 2011). The use of wireless headsets for recording EEG may have increased the noise in the EEG signal, and affected the backbone computations. However, recent studies have shown similar performance as standard EEG hardware (Badcock et al., 2013; David Hairston et al., 2014; Schiatti et al., 2016), making wireless EEG recordings ideal for research in resource-poor settings (McKenzie et al., 2017). Lastly, an additional limitation is that history taking might be affected by a recall bias; in as much patient records (i.e. dates of births) in Nigeria are not stored like they are in modern Western countries. This bias will have increased the noise-level in the regression analysis.

Since both electroencephalography and the use of network backbones reduce data density (i.e. does not take into account peripheral functional network elements), future research should also use, or combine electroencephalography with, other neuroimaging techniques such as functional MRI and magnetoencephalography, which allows mapping of different functional aspects of the brain. Furthermore, in addition to the application of MST analysis, future studies should use other network analysis techniques such as Bayesian exponential random graph models (Caimo and Friel, 2011; Sinke et al., 2016), mixed-effect models (Simpson and Laurienti, 2015) and Gibbs distribution models (La Rosa et al., 2016), which also enable unbiased comparison of networks differing in size and density. This might further elucidate the role of specific brain areas in functional backbone shifts in normal and sensory lacking conditions, which may improve our understanding of neuroplasticity occurring after auditory and other types of sensory deprivation.

### Conclusion

We were able to detect functional network backbone differences in eyes-closed and eyes-open conditions as well as larger shifts in some functional backbone characteristics in deaf as compared to controls. Furthermore, differences were seen between different forms of deafness, and our study demonstrated subtle functional network backbone changes and differences with increasing experience of American Sign Language. Our results provide original insights into the organization of functional brain networks derived from electroencephalography data, both in deaf and healthy people. This might help to better understand functional brain connectivity with lack of auditory input and the development of sign language specifically, as well as cross-modal plasticity due to lack of sensory input in general. Our results further underpin the notion of brain-wide neuroplastic capacities and global network reorganization in the cortex of deaf people. The distinct reorganization patterns between different types of deafness informs on the importance of underlying etiology and staging of the auditory damage in the process of reorganization. The link between the functional network backbone characteristics and acquired sign language experience reflects ongoing brain adaptation in people with hearing disabilities.

## Contributors

MRTS, JWB, EvD and WMO were involved in designing of the study. JWB, MRTS, FvdM, JN and WMO were involved in acquisition of the data. JWB, MRTS and WMO performed the data analysis. JWB, MRTS, FvdM, JN, RMD, EvD and WMO interpreted the data and wrote the manuscript.

